# Biodiversity and Conservation Priority Setting for the Vascular Flora of New Guinea

**DOI:** 10.1101/2022.12.22.521638

**Authors:** Michael Gerard Smith, Felix Forest, James Rosindell

## Abstract

**Aims:** New Guinea is one of the world’s most floristically diverse islands, but its plant collection records are very uneven. We aim to identify which areas have the highest diversity of vascular plant genera, and which areas have the highest deforestation risk. Combining these findings we highlight priority regions for research and conservation.

**Location:** New Guinea

**Time period:** 1900–2021

**Taxa studied:** Tracheophyta (Vascular plants)

**Methods:** We obtained collection records and environmental variables and prepared a cost-distance map of New Guinea to indicate accessibility. We modelled the joint distribution of 1,156 genera with the ‘Hmsc’ package in R to predict biodiversity across space, accounting for collection bias. We combined these results with a genus-level phylogenetic tree to predict phylogenetic endemism. We then modelled deforestation risk with the ‘R-INLA’ package, using forest clearance data and variables including cost-distance. We compared actual and predicted deforestation, and made predictions for 2021–25. Finally, we developed a combined measure of predicted biodiversity plus deforestation risk.

**Results:** A mean Spearman’s rank correlation of 0.462 was obtained on five-fold cross-validation of the genus biodiversity model; bias-correction shifted the predicted distribution of biodiversity towards western New Guinea, but had less effect on estimates of phylogenetic endemism.

Predictions of relative deforestation probability were accurate over 5 and 10 years (Spearman values 0.66 and 0.71). We postulate a ‘deforestation debt’ to explain the persistence in accuracy. Over time, the areas which survive early deforestation gradually become more rewarding targets and the proportion of at-risk forest lost to clearance accumulates.

**Main conclusions:** We present a method for rapid assessment of biodiversity and deforestation risk in data-deficient tropical forest regions. Areas of high predicted biodiversity such as the Merauke and Jayapura lowlands are at high near-term risk from commercial deforestation, requiring urgent interventions to record and preserve threatened species.

## 1. Introduction

As many as one million plant and animal species face extinction because of human activities (IBPES, 2019). While some modern extinctions are relatively well documented, no global analysis until 2019 included plants. Current plant extinction rates are up to 500 times the background rate (Humphreys et al., 2019) and as many as 39% of assessed plant species are threatened with extinction (Nic Lughadha et al., 2020). The biome with the most threatened species is overwhelmingly tropical rainforest, where the greatest threat is anthropogenic habitat conversion (Brummitt et al., 2015).

Conserving plant biodiversity should be a data-driven task (Kindsvater et al., 2018). The 2010 Convention on Biological Diversity Global Strategy for Plant Conservation aims to protect 75% of known threatened plant species, yet barely 10% of plant species are on the IUCN Red List, and many of these are classed as Data Deficient (Pelletier et al., 2018). When assessing priorities for conservation, we aim to identify areas which harbour either rare species, or high levels of species richness or genetic diversity (Forest et al., 2015). We may then predict which of those areas are most vulnerable to disturbance, to focus our protection and management efforts. Many tropical regions, however, are rich in diversity but poor in data (Brummitt et al., 2021).

Here we focus on New Guinea, the world’s most floristically diverse island (Cámara-Leret et al., 2020). Habitats in New Guinea remain more intact than, for example, in Borneo, but plant collection records are very uneven and national boundaries obscure efforts to study the region as a whole. Species distribution models (SDM) that predict distribution ranges using available data offer opportunities to bypass these limitations. Use of such SDMs, however, requires careful handling of sampling bias arising from site accessibility, especially when using ‘presence-only’ methods (Fernández & Nakamura, 2015). Although sample size and choice of modelling method may be as important as spatial bias (Gaul et al., 2020), here we investigate the effects of bias covariate correction (Chauvier et al., 2021; Warton et al., 2013), using a novel cost-distance map as a proxy for accessibility to collectors.

Species are not the only units useful in modelling (Smith et al., 2019). Where records are scarce and most species are rarely recorded, many species will have too few records to make a useful contribution to an SDM (van Proosdij et al., 2016). To use only more common species, however, would produce biased diversity estimates, unrelated to endemism and focused on generalists (Banks-Leite et al., 2014). Rather than modelling genera separately and pooling the results, we model overall generic richness with joint distribution models (Norberg et al., 2019), which help incorporate unmeasured environmental variables. Although bias covariate correction has been investigated using sets of cost and/or distance variables (Bonnet-Lebrun et al., 2020; El-Gabbas & Dormann, 2018), to the best of our knowledge no such study has yet employed a spatial joint SDM. It is also natural to model deforestation spatially, as forest clearance can be ‘contagious’ (Rosa et al., 2013). There are ‘hotspots’ of deforestation in areas where commercial oil palm plantations (The TreeMap, 2021) have been identified.

New Guinea has “arguably the world’s most complex geotectonic history” (Toussaint et al., 2014). It is divided between independent Papua New Guinea (PNG) and two Indonesian provinces (Papua and West Papua, collectively ‘Indonesian Papua’) (see Figure 1). Geologically isolated regions include the Bird’s Head peninsula in Indonesian Papua, and in PNG, the islands of New Britain and New Ireland and the mountainous Huon peninsula (Baldwin et al., 2012). An inventory of 13,634 identified species, of which 68% are endemics, is available (Cámara-Leret et al., 2020), but the distribution and abundance of these species are largely unknown. The GBIF database (GBIF.org, 2021) contains 38,475 vascular plant records in New Guinea, identified to species level and geolocated; 78% of the species identified in this dataset have five specimens or less, as do 73% of the recorded species in LAE, the national herbarium of PNG (Conn et al., 2006). The median year of first description for endemic species is 1927, and the geolocation data of earlier specimens was usually poor. Of the listed endemic species, 53% have been found only in PNG and 24% only in Indonesian Papua; however, a large proportion of collections has been made in PNG (Cámara-Leret et al., 2020).

**Figure 1.**
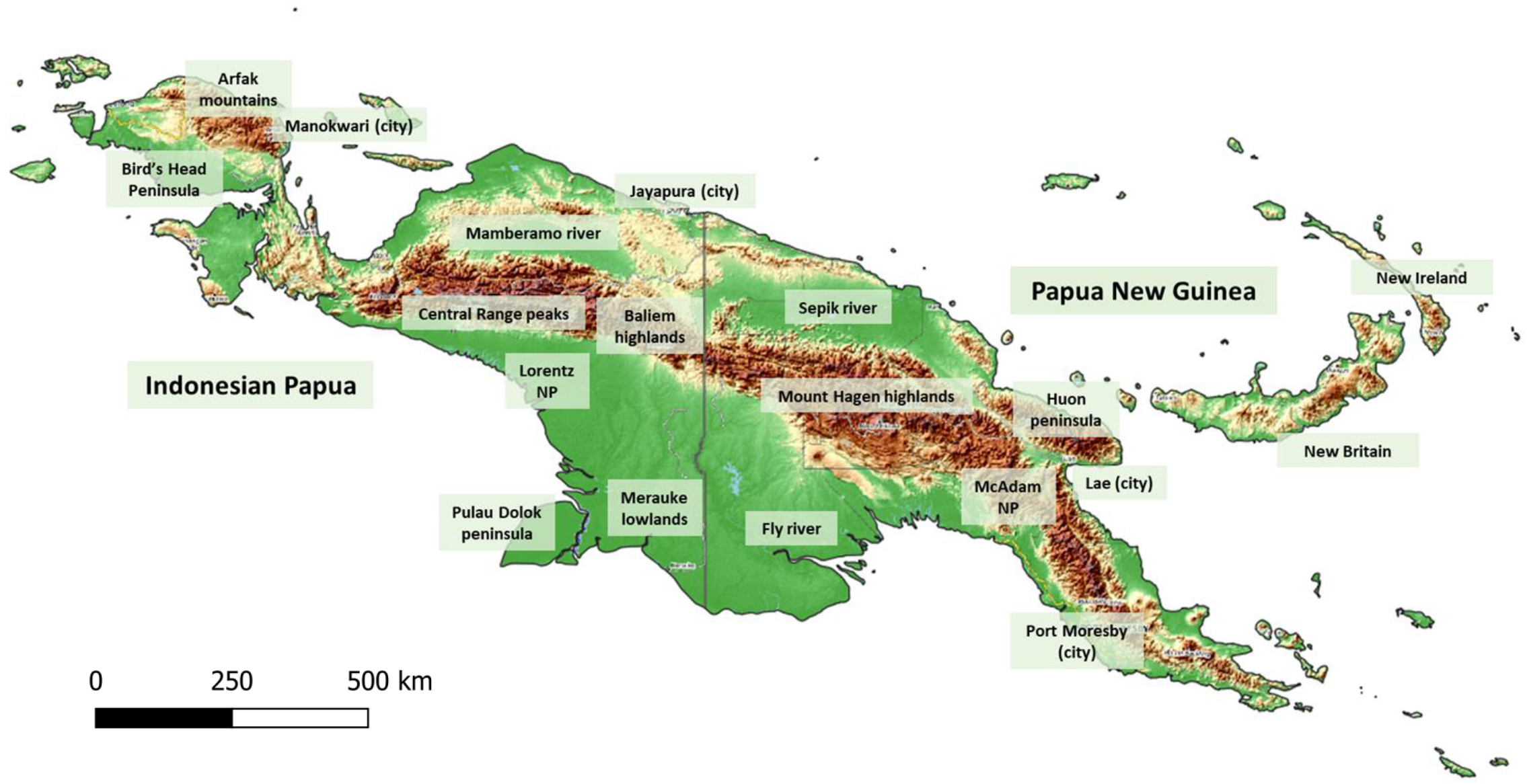
Map of New Guinea, showing the major mountain ranges, islands and peninsulas, river/wetland areas, and population centres. The area covered by this study includes New Britain and New Ireland but excludes Bougainville (effectively independent and closer, floristically, to the neighbouring Solomon Islands).

The population of PNG exceeds 8.5 million people, with widespread agriculture throughout the highlands, but that of Indonesian Papua, a region of equivalent extent, remains below 5.5 million, with inland populations concentrated in the Baliem valley. Primary forest extent in Indonesian Papua was 86.9% in 2000 and remained at 86.2% in 2012, a period during which Sumatra and Indonesian Borneo had lost 17.6% and 7.9% of primary forest, respectively (Margono et al., 2014). Although the situation in PNG is less clear, it has been reported that 63.4% of PNG is forested, and that 91.2% of this is primary forest (Mongabay.com, 2021).

Deforestation rates and mechanisms differ across New Guinea. A 3.7% decrease in tree cover was reported in PNG from 2000–2020, with a 2.4% loss of primary humid forest. Figures for Indonesian Papua show a 2.3% decrease in tree cover from 2000–2020, with a 1.8% loss of primary humid forest (Global Forest Watch, 2021). In PNG, 63% of current forest loss is non-commercial (United Nations Framework Convention on Climate Change, 2019) and the figure for Indonesian Papua is 55% (Gaveau et al., 2021). Forest loss in New Guinea as a whole peaked in 2015 and has since fallen to a level below that of 2012 (Hansen/UMD/Google/USGS/NASA, 2021); the rate of deforestation in PNG may have fallen to 0.5%/year by 2015 (United Nations Framework Convention on Climate Change, 2019). Future threats will arise from the still-incomplete Trans-Papua Highway, which will improve access to plantations in Indonesian Papua (Gaveau et al., 2021).

In the near future, loss of habitat in New Guinea will depend on population pressure, the political and economic situation, and the progress of oil palm and pulpwood concessions. Here we aim to predict relative deforestation risk over 5–10 years, using publicly available data. We use a cost-distance map developed in QGIS (QGIS.org, 2021) as a proxy for accessibility but dispense with the need for time-consuming visual calibration of remote sensing images.

We will predict which regions have higher biodiversity in terms of genera, identify areas most vulnerable to deforestation, and highlight areas combining high diversity and high risk as priority targets for research and conservation. The ‘Hmsc’ package in R (Tikhonov et al., 2020) allows us to include cost-distance, together with occurrence records and environmental covariates, in a spatially explicit SDM; the ‘R-INLA’ package (Blangiardo et al., 2013) allows cost-distance to be used in a spatial model of deforestation. Using this dual approach we highlight areas combining high predicted biodiversity and high relative risk of deforestation (Molotoks et al., 2017).

## 2. Methods

We downloaded geolocated species collection records, deforestation maps, and a suite of predictor datasets, and processed them for subsequent regression analyses. For use as an additional predictor, we developed a cost-distance map of New Guinea, which provided a quantification of the accessibility of a given area to plant collectors. We modelled biodiversity and the risk of deforestation, and calculated phylogenetic endemism (PE); we validated the biodiversity models with multiple (x40) cross-validation exercises. We then combined the results from the models to identify important areas combining both high generic diversity and high risk of deforestation.

### 2.1 Collection records

We obtained collection records, with permission (Conn et al., 2006), from the Papua New Guinea Forest Research Institute herbarium (LAE) and also from GBIF (GBIF.org, 2021); records did not overlap, as the LAE records do not appear in GBIF. Other relevant herbaria (including the Herbarium Bogoriense in Jakarta, the Singapore Botanical Gardens, and the Naturalis Institute of Leiden) have few or none of their New Guinea specimen records available for download.

In total, 18,796 georeferenced records from GBIF and 115,386 from LAE had been identified at species level and assigned to 13,445 species based on the recent checklist for the island (Cámara-Leret et al., 2020). We mapped these records in QGIS (QGIS.org, 2021) and excluded those mislocated in the sea. We excluded records from LAE if either latitude or longitude was recorded with less than 1/4 arc-minute of accuracy. Records from GBIF, which uses decimal degrees, were excluded if either measurement was stated to less than 0.05 degree accuracy (or if it appeared that a whole degree had originally been measured). As GBIF records come with an estimate of location inaccuracy, we excluded those with estimated imprecision >3,000m; this information was not available for the LAE records. Finally, 27,367 records, representing 1,156 genera and 6,180 species, were used; 40% of these were from GBIF and 60% from LAE. We excluded duplicate species-site records.

### 2.2 Deforestation and environmental variables

We calculated standardized PCA values from rasters of 101 continuous environmental variables (Table S2), mapped across the whole region. Variables considered included Worldclim (WorldClim, 2021) and ENVIREM (Title & Bemmels, 2018) climate variables, soil variables from the International Soil Reference and Information Centre (Poggio et al., 2021), and derived measures of soil quality from the UN FAO (Food and Agriculture Organization of the United Nations, 2021). An elevation map was used (United States Geographic Service, 2021) in case it could supply information missing from datasets whose data were interpolated from scattered meteorological stations (Fletcher et al., 2016). We used three MODIS satellite datasets from 2003 (Myneni, 2015): fraction of photosynthetically active radiation absorbed by green vegetation, leaf area index, and enhanced vegetation index.

We obtained population data from the WorldPop2020 database, yearly from 2000–2020 (University of Southampton, 2021). We also obtained current protected area information (United Nations Environment Programme, 2021), but historical data is embargoed. The current Indonesian land legal classification was available, but as no equivalent dataset was available for PNG, we excluded this from the analysis.

### 2.3 Cost-distance analysis

We built a cost-distance map in QGIS (QGIS.org, 2021) (see Supporting Information). We used corrected digital elevation model values in conjunction with a land-system map (Saxon & Sheppard, 2010) that incorporates land categories from the Regional Physical Planning Project for Transmigration in Indonesia, plus the land units of the Papua New Guinea Resource Information System. At a resolution of 1km^2^, we derived a normalized mean slope raster from the elevation map; this was multiplied by a cost raster which categorized the difficulty of crossing different types of terrain in New Guinea, informed by personal experience (Supporting Information Table S1).

We prepared a raster to map the minimum cost-distance from towns/ferry-ports/airstrips (Humanitarian OpenStreetMap Team, 2021) to every part of the region. As ferry services in New Guinea are often modern and reliable, whereas roads may be less so, we assigned sea and roads equal cost value and extended the cost-distance map to 5km offshore. The border between PNG and Indonesia Papua, which until recently was officially impermeable, was set to the maximum cost value.

### 2.5 Biodiversity predictions

To investigate the applicability of various SDM techniques, we re-purposed a workflow from a recent study which examined the performance of a range of SDMs (Norberg et al., 2019). Within the context of a priority-setting framework, relative ranking of biodiversity is an appropriate measure (Guillera-Arroita et al., 2015) and so, of the possible predictive performance measures, we focused on the Spearman rank correlation between observed generic richness and predicted values. We used QGIS to prepare a map of the 180 plant species recorded in ≥30 New Guinea grid squares of 25km^2^ size; we split the grid squares randomly into 50% training and 50% validation units, and species found in only one of the two sets were discarded. Having tested model outputs under two-fold cross-validation, we identified the ‘Hmsc’ package (Tikhonov et al., 2020) as the most successful model.

The ‘Hmsc’ package allows joint SDMs to be developed with an independent spatial term. Our model used five PCA axes and the cost-distance variable; we compared it to a pooled ensemble of single-genus models using ‘Hmsc’. We jointly modelled the presence of 1,156 genera, whose recorded distribution covered 313 of the available 549 grid squares of size 50km^2^. Grid centroids were used for the coordinates of the spatial random effect. To fit each model, we used ‘Hmsc’ to run two MCMC chains of 1,000 samples with thinning x5 and therefore 10,000 iterations, discarding the first 2,500. The trace plots were checked for satisfactory convergence, as were the Beta-coefficients of the fitted model.

We tested models with 40 trials of five-fold cross-validation; 5-folds were prepared by random subsetting of 313 grid squares, with 2,000 sets of predictions sampled in each trial from the posterior predictive distributions of the five partial models. Next, we predicted biodiversity predictions across all grid squares in New Guinea, using a model fitted with data from the 313 grid squares holding collection records. Again, 2,000 samples were drawn to make predictions. Finally, we re-fitted the model with the cost-distance variable set to zero, to correct for sampling bias (Warton et al., 2013) and further predictions were extracted.

### 2.6 Phylogenetic endemism

Using the R package ‘V.Phylomaker’ (Jin & Qian, 2019), we built an ultrametric phylogenetic tree for all species found in New Guinea of the genera used in the biodiversity analyses. The tree was then pruned to a single species per genus to obtain a genus-level tree. We used the BIODIVERSE spatial analysis application (Laffan et al., 2010) to calculate unweighted phylogenetic endemism (PE) across New Guinea using the genus-level tree (BIODIVERSE subroutine: calc_pe_single, cell size 50km^2^ (Rosauer et al., 2009)).

We first calculated PE from the original collection record dataset and then from the modelled biodiversity predictions across New Guinea (both the original and bias-controlled model). BIODIVERSE allows the use of fractional species presence probabilities; however, we excluded modelled probability values for genus presence if they were more than two standard deviations below 0.5.

### 2.7 Deforestation risk

We calculated deforestation using the Global Forest Watch 2000–2020 global dataset (Hansen/UMD/Google/USGS/NASA, 2021) which records the earliest year of clearance. We did not consider reforestation, as the main causes (acacia for pulpwood, and oil palm) are unlikely to restore lost biodiversity. We divided New Guinea into 10km^2^ grid squares and investigated regression models, with the response variable being the number of deforested 30m^2^ pixels in a 10km^2^ area (excluding any parts of the grid squares covering sea). Of the 10,066 grid squares, 215 (2.1%) changed their majority vegetation cover over 20 years, 139 of which (65%) were originally majority forest. Given the low rate of change, we modelled vegetation type with a single dataset from 2000 (ESA/MEDIAS-France-POSTEL, 2021).

We developed a model in ‘R-INLA’ (Blangiardo et al., 2013), hypothesizing a zero-inflated negative binomial distribution of cumulative yearly forest clearance. A Besag-York-Mollie spatial random effect was incorporated, applied to a neighbourhood lattice graph of the grid squares (Blangiardo et al., 2013). We made predictions over 5 and 10 years using models fitted with partially withheld data; to judge relative predictive accuracy, we compared these to the actual data using Spearman’s rank correlation.

As predictor variables, we used time (year from 2000), mean cost-distance, mean elevation (slope was already included in the cost-distance variable), the proportion of a grid square falling within an officially protected area, and the majority vegetation type. We also considered estimated population density. A model fitted with the entire dataset was extrapolated to make predictions for the period 2021–25.

### 2.8 Biodiversity and risk

Using QGIS, we merged the predicted deforestation for 2021–2025 from 10km^2^ into 50km^2^ grid squares, and took the logarithm of predicted values across New Guinea for both deforestation and biodiversity. We normalized each set of values to a range of 0–1; the results were added, giving a composite measure of biodiversity and deforestation risk per grid square, and mapped by decile. We produced maps of combined risk using both original and bias-controlled diversity predictions.

## 3. Results

### 3.1 Cost-distance analysis

Only 11% of collection records were from Indonesian Papua, and 1,118 of 1,834 25km^2^ grid squares across New Guinea had no records in the data sets we used. Having hypothesized that ease of access affects both collection records and deforestation risk, we developed cost-distance calculations for use in modelling both (see Supporting Information Figure 1).

In Indonesian Papua especially (see Supporting Information Figure 2), collection sites are associated with convenient locations, e.g. the cities of Jayapura and Manokwari, the road from the coast to the giant Freeport mine in the highest peaks of the Central Range, and the populous Baliem Valley highlands.

**Figure 2.**
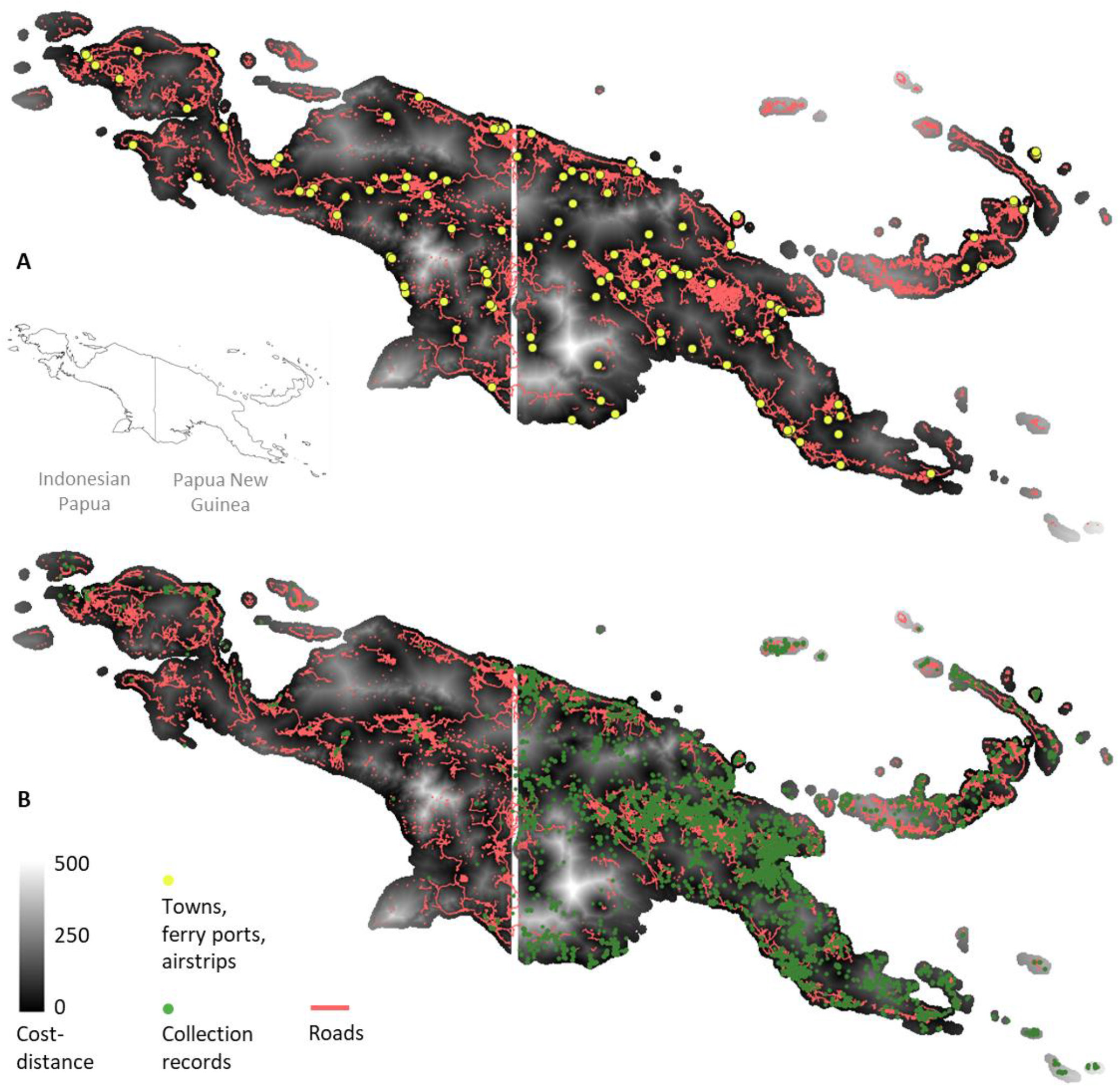
Cost-distance map. Scale from lowest to highest access cost. A, cost-distance with roads and towns/ferry ports/airstrips. B, cost-distance with roads and collection sites. The international boundary was set to the maximum cost-distance value.

The median cost-distance of collection points was 61 (mean 79, see Supporting Information Figure S3), which is lower than the average value of the cost-distance map (median 102, mean 121, Wilcoxson rank-sum test p=2.2e-16). Nevertheless, there is little correlation between mean cost-distance per 50km^2^ grid square and number of collection records (see Figure S4). Many areas with low cost-distance are, perhaps by chance or because of previous clearance, also lacking in records.

### 3.3 Biodiversity predictions

The fitted ‘Hmsc’ model had mean *AUC* across grid squares of 93%. Using five-fold cross-validation to compare predicted to observed generic richness, a mean Spearman value for the models of mean 0.462 (*SD* = 0.01), standard deviation of Spearman values 0.036 (*SD* = 0.001) was obtained across the 40 trials of five-fold permutations. Dropping the spatial term in the regression reduced the Spearman value by 48%; keeping the spatial term, but modelling genera individually and pooling the results, reduced the Spearman value by 42%. The addition of squared predictor variables to the model provided no extra benefit.

We predicted higher generic biodiversity in montane regions, especially the Central Range and east New Britain (see Figure 3). Collection records showed little variation in biodiversity across the Bird’s Head peninsula, but the model predictions suggested higher biodiversity in the peninsula’s mountains (the Arfak range). Predicted biodiversity was generally higher in PNG than in Indonesian Papua, where data is scarce. However, re-fitting the model with the cost-distance variable set to zero showed a pronounced increase in predicted biodiversity between 133°–143° longitude (mostly Indonesian Papua), and a corresponding decrease in PNG between 143°– 151° longitude. We predicted high biodiversity the southern coast and the drier lowland areas of Indonesian Papua.

**Figure 3.**
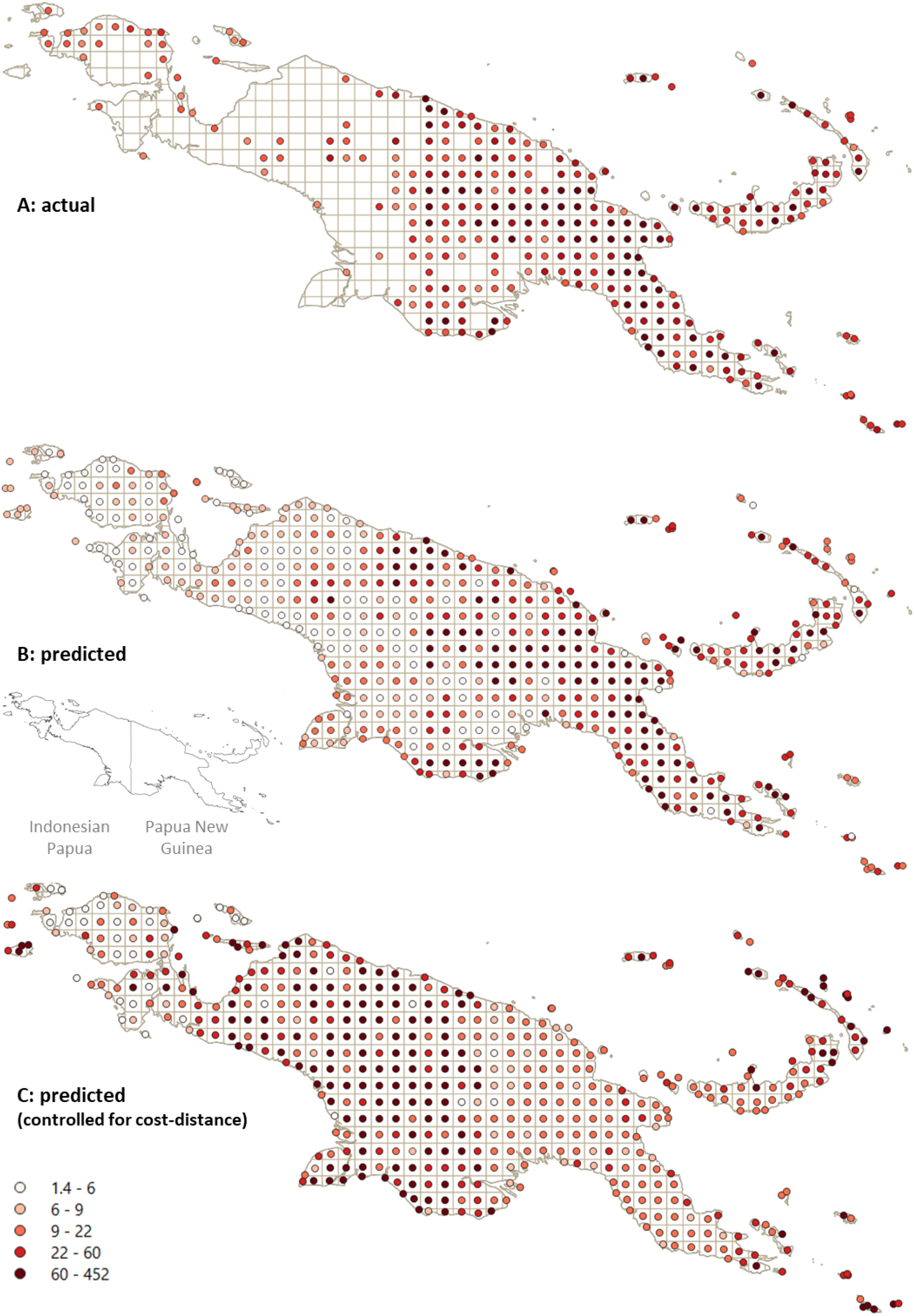
Actual and predicted biodiversity (number of genera). A, actual records, sampled 50km^2^ grid squares only. B, predictions with ‘Hmsc’ model using 5 PCA axes plus the cost-distance variable. Biodiversity is predicted to be higher in montane regions and in drier lowlands. C, biodiversity predicted from the fitted model with the cost-distance variable controlled. Inset shows international boundary.

### 3.4 Phylogenetic endemism

The actual collection records produced PE values strongly skewed towards areas of higher collection effort, although we saw a trend towards higher PE in montane regions. The modelled PE values showed a concentration of PE in areas also modelled as having high biodiversity (see Figure 4; for PE values based on the model before bias-correction, see Supporting Information Figure S5). Predicted PE before and after bias-correction was comparable (Spearman’s rank correlation = 0.72). Predicted PE was low in the Bird’s Head peninsula and in central mainland PNG, and high in central mainland Indonesian Papua.

**Figure 4.**
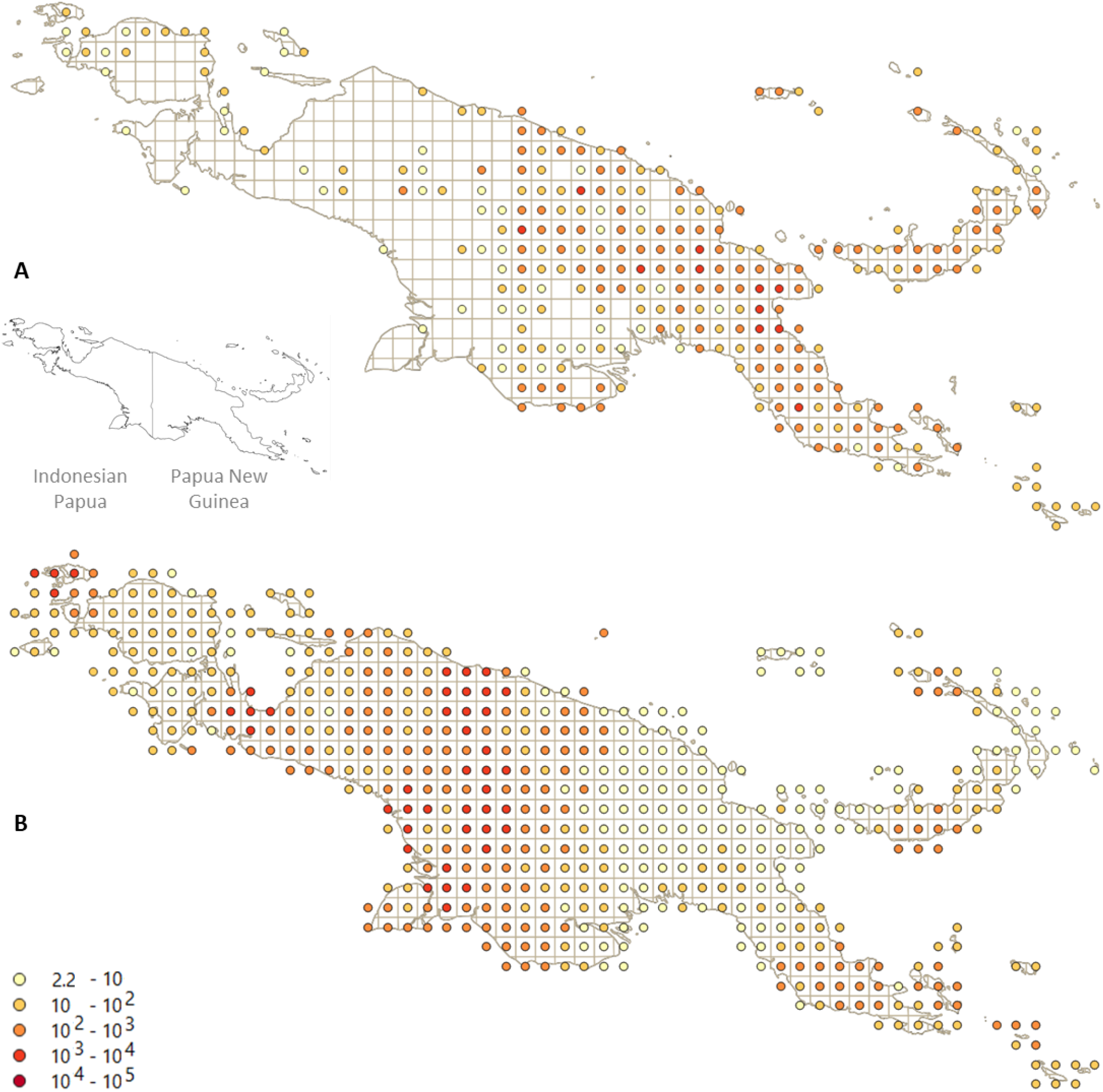
Phylogenetic endemism (log scale, depicted by quintile). A, estimated from actual records. B, predicted from ‘Hmsc’ model (5 PCA variables, cost-distance variable set to zero). Note that as PE is calculated in a neighbourhood-based lattice of 50x50km squares, values are not displayed as ‘snapped’ to grid centroids. Inset shows international boundary.

### 3.5 Deforestation risk

Using a model fitted with data from 2000–2015 to make predictions for 2016–2020, the Spearman rank correlation between actual and predicted deforestation over the latter period, by grid square, was 0.66 (to compare this to a null value, 1,000 random permutations gave a correlation of -0.0001, *SD* 0.0044). The median predicted deforested pixels per grid was 1,906 (mean 7,628) compared with actual values of 1,158 (mean 2,917), giving the predicted distribution a longer tail than the actual distribution. The mean absolute error of the prediction was 5,207.

The same model fitted with data from 2000–2010 gave predictions for 2011–2020 with a relatively high Spearman value of 0.71. For future use, the model fitted with data from 2000–2020 (see Table 1 for coefficients of the fixed effects) was used to make predictions for the years 2021–25 (see Figure 6).

**Table 1.** Fixed effects coefficients of the deforestation regression model. Low cost-distance, low elevation, situation outside protected areas, and more recent year predict more deforestation, as do vegetation types with mixed forest/cultivated cover in 2000. Less deforestation is predicted in areas with vegetation types susceptible to seasonal or regular flooding.

|  | Mean | SD | CI: 0.025 quantile | CI: 0.975 quantile |
| --- | --- | --- | --- | --- |
| (Intercept) | 4.354 | 0.075 | 4.207 | 4.500 |
| Cost-distance (z-scaled) | -0.295 | 0.007 | -0.309 | -0.282 |
| Elevation (z-scaled) | -0.299 | 0.007 | -0.244 | -0.215 |
| Proportion within protected area | -0.190 | 0.024 | -0.237 | -0.143 |
| Vegetation: mostly cropland (>50%), some natural | 0.152 | 0.083 | -0.012 | 0.316 |
| Vegetation: mosaic, mostly natural (>50%), some cropland | 0.086 | 0.105 | -0.120 | 0.291 |
| Vegetation: tree cover >15% | -0.162 | 0.073 | -0.305 | -0.019 |
| Mosaic herbaceous cover, tree and shrub | -1.363 | 0.093 | -1.544 | -1.181 |
| Tree cover, flooded, fresh/brackish water | -1.435 | 0.080 | -1.592 | -1.279 |
| Tree cover, flooded, saline water | -2.080 | 0.080 | -2.236 | -1.923 |
| Year (from 2000) | 0.178 | 0.001 | 0.177 | 0.179 |

Predicted and actual deforestation (see Figure 5) was concentrated in lowland areas free from seasonal or permanent flooding, and in mid-altitude highland valleys. These areas are more populous, and the former are suitable for commercial plantations.

**Figure 5.**
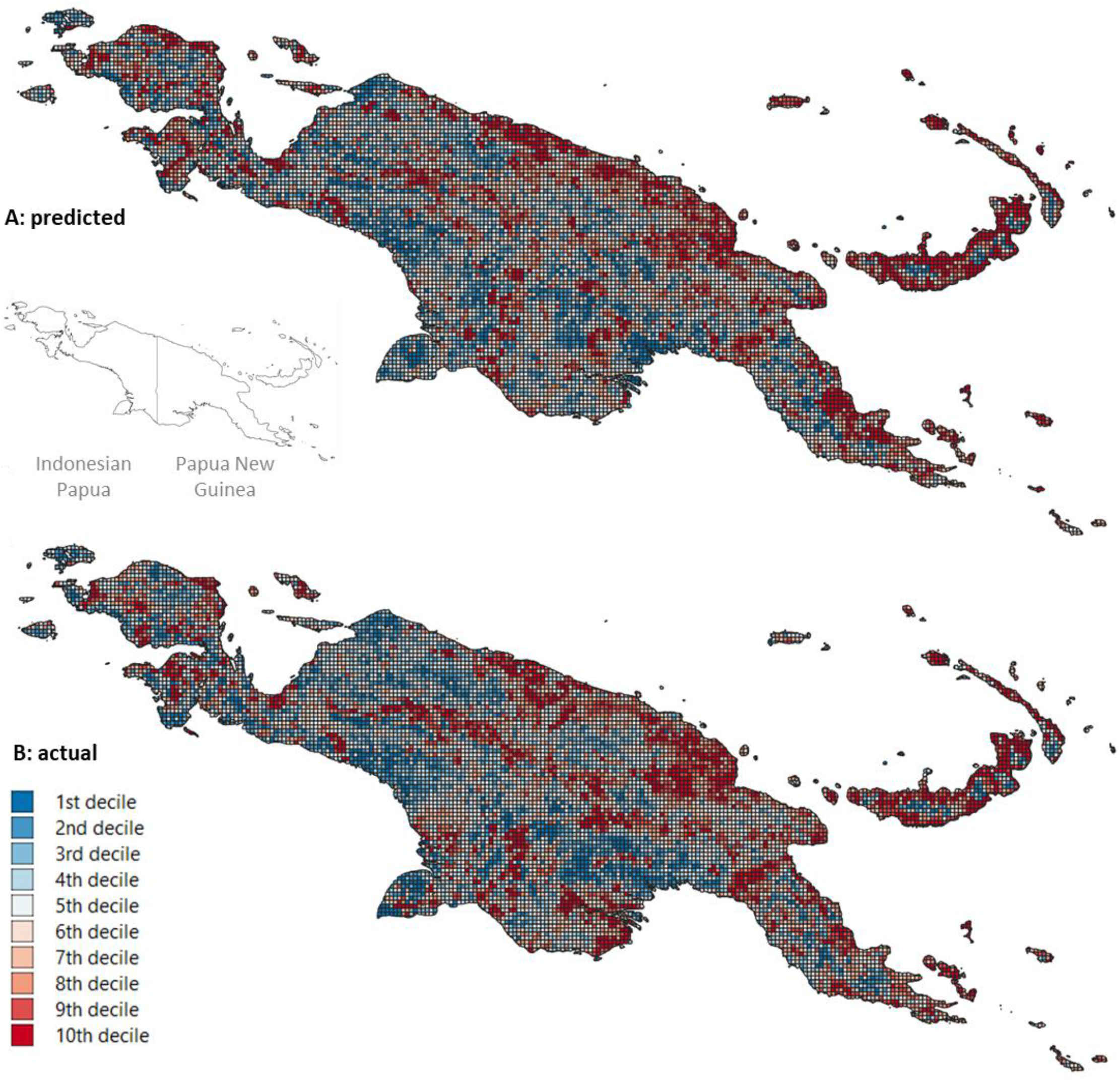
Predicted versus actual deforestation; relative deforestation by decile. A, relative deforestation risk over 2016–20 predicted from 2000–2015 data; B, actual deforestation 2016–2020. Inset shows international boundary.

Population density, mostly very low, increased in many areas from 2000–2020. Population density per 10km^2^ grid square changed by a factor of median 1.91 (mean 3.59, *SD* 4.76). As a regression predictor, however, it had no additional effect.

We used various non-parametric estimation methods to investigate potentially complex relationships between the variables and the response. However, adding smoothing terms in the linear predictor (by binning the predictor variables into groups by range, and fitting penalized splines to the data according to a random walk model) did not improve model fit (*WAIC*, Watanabe-Akaike information criterion) or predictive accuracy. We applied the model to 5km^2^ and 2km^2^ grids, with no improvement in predictive power.

### 3.6 Biodiversity and risk

Areas in PNG with a combination of high biodiversity and high risk included not only the central highlands (where population pressure is high) and eastern New Britain (which has seen rapid commercial deforestation), but also a number of islands and lowland coastal areas suitable for plantation (see Figure 7). In Indonesian Papua, lowland areas with commercial concessions within reach of the nascent Trans-Papuan Highway – including the Merauke region and the region around Jayapura – combine high diversity and high risk.

## 4. Discussion

We applied recently developed techniques in statistical computation to estimate vascular plant diversity across New Guinea and the regional variation in the risk of deforestation, the main cause of non-natural habitat loss. We assembled a wide variety of predictor datasets for use in our modelling approach. Some of these datasets were highly collinear with others, some relied on interpolation between data recording stations, and not all fitted the land/sea boundaries of New Guinea precisely. Nevertheless, they could be reduced to efficient predictor sets by PCA, as demonstrated by model cross-validation. We developed an original measure of cost-distance as a proxy for accessibility. Despite some subjectivity, it had a demonstrable influence on deforestation risk (see Table 1), and affected the measurement of biodiversity via the distribution of collection records (see Figure 2). The relationship between high cost-distance and number of collection records was significant (see Additional Information Figure S3), though some ‘difficult’ areas, such as mountain peaks with alpine flora or swamps rich in pitcher plants (Nepenthaceae), may attract rather than deter exploration.

The low biodiversity predicted for the Bird’s Head peninsula (see Figure 3) may indicate that modelling cannot overcome the lack of data in this outlying region (the lowest quantiles of diversity are implausibly low, even when controlled for cost-distance). Conversely, the model predictions for areas near Lae and Port Moresby seem more likely than the higher biodiversity estimates from actual collection records, which may be a product of easier access and intensive collecting. The ongoing development of a network of forest plots in PNG (Center for Tropical Forest Science, 2021) could assist ‘ground-truthing’ the results obtained from these models.

PE patterns in New Guinea deserve more attention, given its status as a hotspot for ancient taxa such as Cyatheaceae and Dicksoniaceae. Model predictions of PE distribution (regardless of bias correction) were very different from the PE compiled from empirical collection records only (see Figure 4, Additional Information Figure S5). If the large landmass of central New Guinea really contains higher PE than the smaller Bird’s Head or the eastern peninsulas, this may suggest a species-area relationship (if high PE and high species richness are related) and/or the potential for larger mountainous areas to act as refuges during climatic change (Roos et al., 2004). However, PE is predicted to be relatively high in the narrow ‘neck’ of the Bird’s Head and on Waigeo Island.

The deforestation model achieved high accuracy in terms of relative risk. The slightly greater accuracy of 10-year predictions to 5-year predictions was unexpected, but it may be that the predictor variables, over time, outweigh shorter-term stochastic variation in forest clearance. Eventually, areas suitable for clearance which have escaped damage in the short term may end up paying a ‘deforestation debt’ as accessible forests become scarcer and the areas which have survived deforestation over 5 years gradually become more rewarding targets. Over time, the proportion of at-risk forest lost to clearance will accumulate.

Our model underestimated risk in southern coastal areas (see Figure 5), which may be more susceptible to dry-season burning (but also to river flooding, a natural and temporary loss of habitat which may be misleadingly recorded as deforestation in remote sensing maps). The deforestation and biodiversity models, combined, suggested a ‘red alert’ combination of high biodiversity and high risk both in over-populated highlands and in lowland areas threatened by oil palm plantations (see Figure 6A) However, risk may change over time as road construction affects accessibility, and the cost-distance calculation should therefore be updated to keep models relevant.

**Figure 6.**
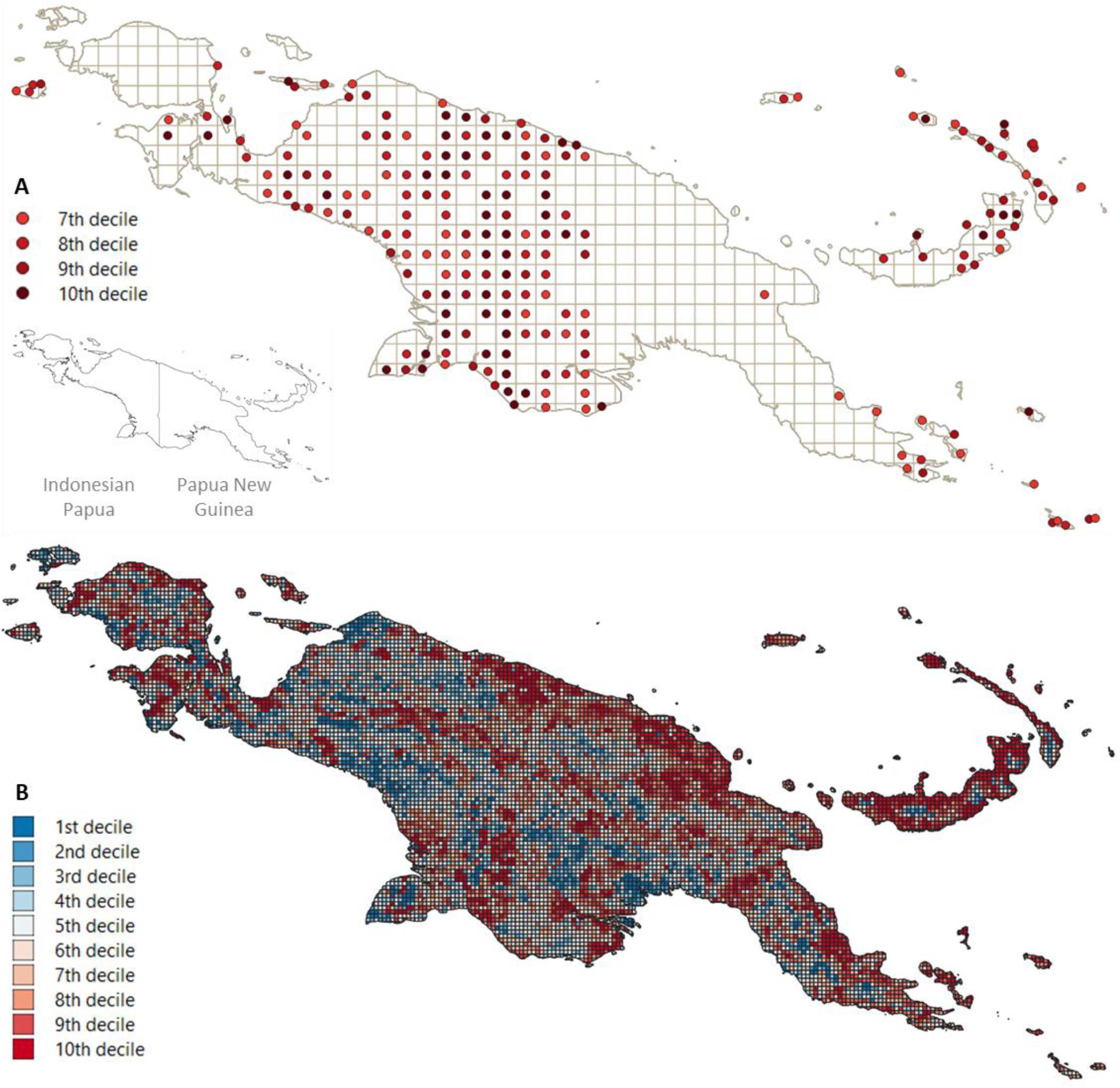
A: Map showing composite measure of predicted biodiversity (bias-controlled model) combined with predicted deforestation (to 2025). Only the four highest deciles are shown (for complete maps – derived from controlled and uncontrolled models – see Supporting Information Figure S4). B: Map showing additional predicted deforestation 2021–25. Inset shows international boundary.

Lists of species from GBIF and LAE could be compared to a checklist of New Guinea plant species (Cámara-Leret et al., 2020), but since the checklist was expert-curated and follows the World Checklist of Vascular Plants, it often diverges from the naming conventions used by GBIF. Detailed review of collection records is needed (Goodwin et al., 2015) to correct misidentification and remove synonyms in genus and species names; this might allow biodiversity estimates to be made based on species as well as genera. Other limitations of the data include imperfect geolocation, uneven efforts towards digitization, and differences in geolocation standards. In future, more herbaria will digitize and georeference their collections, which should improve the balance of records across the two halves of New Guinea.

If collection records are uneven, biodiversity is easier to estimate at a larger spatial scale; the absence of records within a small area may be due to chance, and at a finer grid scale there will be more areas without records. In contrast, deforestation should be modelled at the highest resolution available. Even 10km^2^ squares can include a range of vegetation types, which may have reduced the precision of ‘majority vegetation type’ as a predictor variable. Deforestation data at 30m^2^ resolution were used, but covariate data at this scale are not readily available. Applying the deforestation model to smaller grids resulted in no significant improvement in predictive power; thus it seems that a 10km^2^ grid was sufficient to capture the scale of spatial dependence. Although we assumed that the deforestation dataset is reliable for New Guinea, global forest cover datasets would benefit from local ‘ground-truthing’ to distinguish, for example, between natural flood damage and deforestation.

There was a noticeable geographical difference in biodiversity predictions made from the original versus bias-controlled models, which deserves further investigation. We suggest that biodiversity estimates made on the basis of collection records skewed by accessibility are unreliable and biased towards areas easily reached by road from population centres. Biodiversity predictions made across all New Guinea (including areas where no collection records were available) cannot be tested by cross-validation and are hard to assess on the basis of internal consistency (Newbold et al., 2010); we made predictions with the principal aim of supplying hypotheses to be tested by fieldwork.

Deforestation has been the subject of complex econometric models involving crop yields, farm-gate prices, and farmers’ dependence on forest resources. Little is known about how these factors affect New Guinea, although this has been investigated in Borneo (Schoneveld et al., 2019). Global Forest Watch data has been used to model deforestation in Indonesian Papua (Gaveau et al., 2021). They inspected satellite images to differentiate clearance from natural forest loss, the expansion of roads over time was recorded, and concession maps were pieced together from datasets compiled by Indonesian NGOs. Forest loss was simulated using a cellular automaton method which identified as the best model the one in which the predictors were active concessions, protected area status, and cost-distance from major roads.

Lacking data on concessions in PNG, here we present a model based on publicly-available data of medium-to-high spatial resolution, without the need for intensive visual calibration but with a well-developed cost-distance element; we applied this model to the whole island and it could easily be repurposed for use in other study areas.

Across New Guinea, and especially in the central ranges of Indonesian Papua, mountainous areas are likely to be rich in biodiversity despite the sparsity of the available natural collections from these regions. These areas are at low risk from commercial plantation, but they are threatened by clearance and burning as a side-effect of population growth in the highlands. In PNG, the threat from commercial deforestation is especially acute in lowland New Britain, and in Indonesian Papua, areas at risk include the neck of the Bird’s Head peninsula, the Jayapura region, and the south-east (Merauke regency). More fieldwork is necessary to investigate how closely our estimates of generic biodiversity match reality; meanwhile, efforts to collect seeds and specimens are urgently required in order to record and preserve species which may otherwise disappear without trace.

Continued developments in Bayesian modelling techniques allow increasingly reliable estimation of both biodiversity and deforestation risk for regions such as New Guinea, an island vitally important for plant conservation but with poor data availability. Despite their limitations, these methods are able to cope with the shortage of data and to inform the practical development of conservation priorities, identifying areas combining high diversity and high risk such as the Merauke and Jayapura lowlands (see Figure 6A). The predictive ability of our biodiversity model, under cross-validation, was comparable to trials of similar models (Norberg et al., 2019) made on the basis of tidier data; our deforestation model would allow a rapid assessment of relative risk in similar data-deficient regions. These predictions could be improved with more accurate data sources, but in the case of New Guinea, as with other reservoirs of tropical humid forest, there is no time to lose.

## Data Availability Statement

Code script is available at: https://github.com/gaiteiro/new_guinea

