## Supplementary Data for "Biodiversity and Conservation Priority Setting for the Vascular Flora of New Guinea"

### Supporting Information

#### 1. Code and applications

Code was developed in R (R Core Team, 2012) and Matlab (The Mathworks Inc., 2021) and data was further processed in Microsoft Excel (Microsoft Corporation, 2018). Instructions for installing the current version of the HMSC package can be found at https://github.com/hmsc-r/HMSC, for R-INLA at https://www.r-inla.org/download-install, for V.Phylomaker at https://github.com/jinyizju/V.PhyloMaker, and for BIODIVERSE at https://github.com/shawnlaffan/biodiverse/wiki/Downloads.

#### 2. Supplementary figures and tables

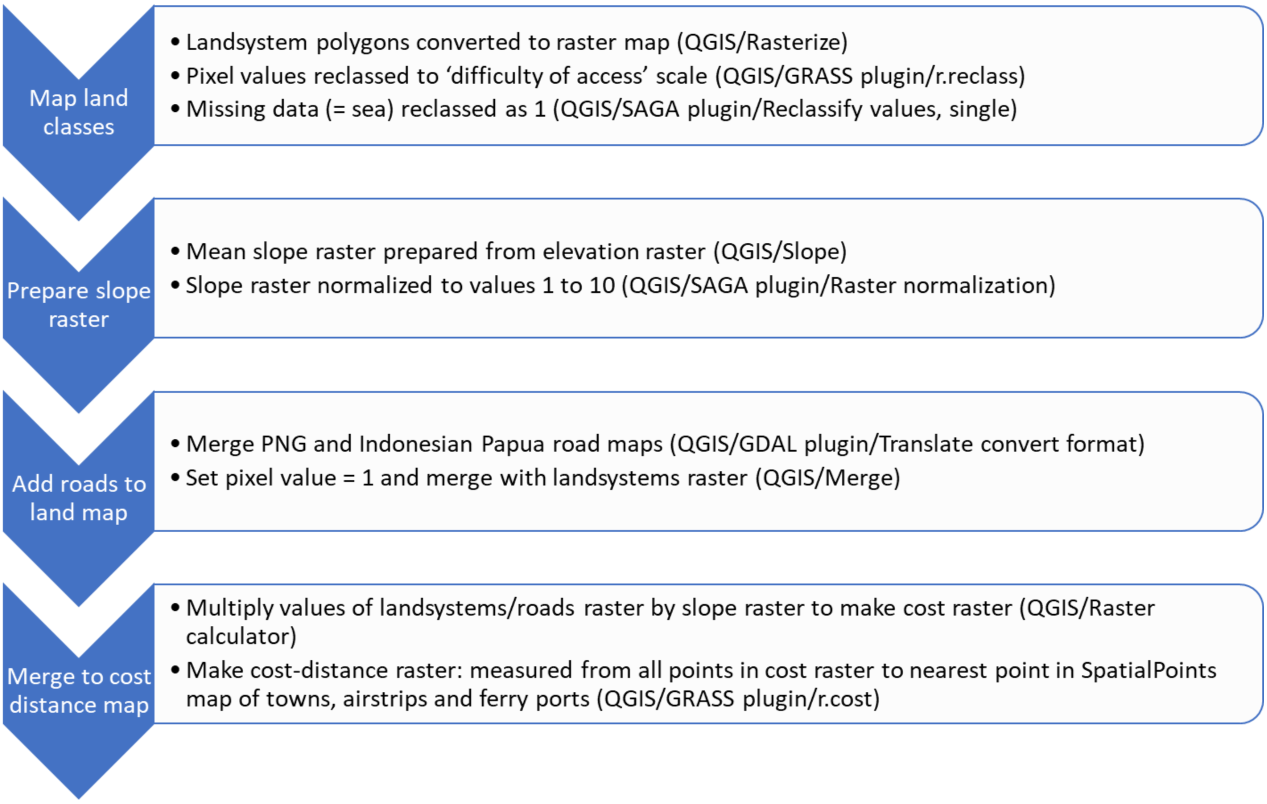


*Figure S1. Flowchart showing development of the cost-distance map. Processing was performed in QGIS (QGIS.org, 2021).*

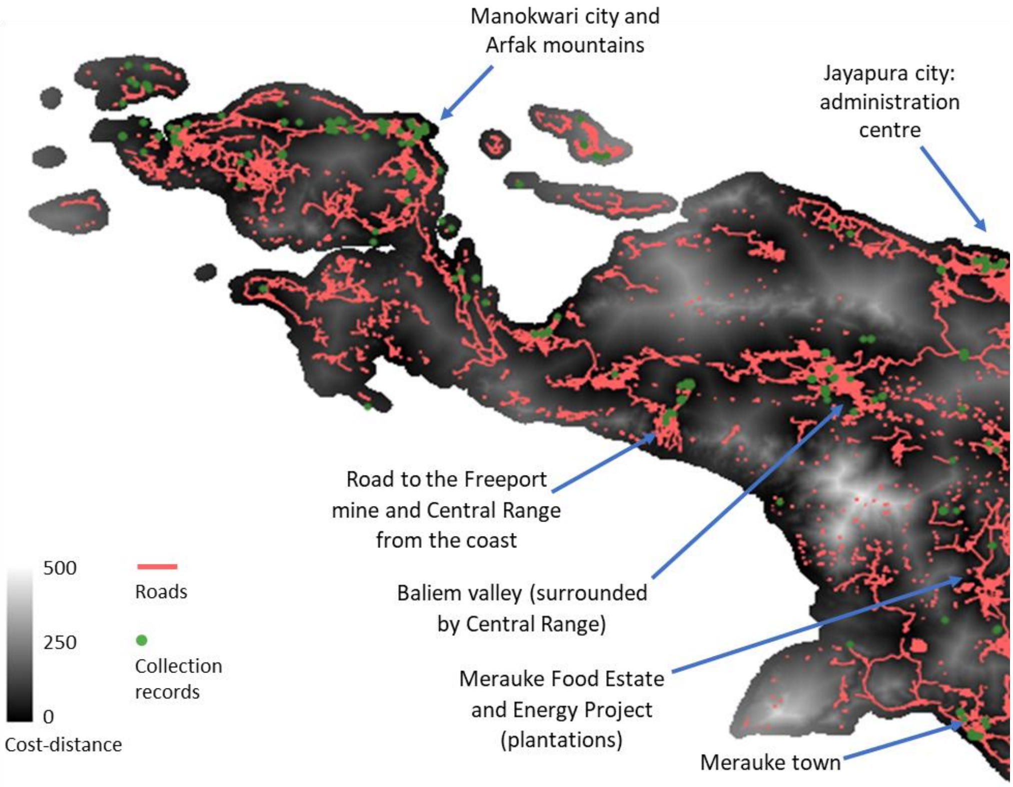


*Figure S2. Cost-distance map of Indonesian Papua. Scale from black (easiest access) to white (most difficult access) with roads shown in pink and plant collection records in green. The Merauke Integrated Food and Energy Estate (MIFEE) is a group of concessions totalling 2.5mHa, projected as a source of food and fuel for the wider Indonesian population but, in practice, mainly consisting of oil-palm plantations owned by investors including LG and Daewoo (Asian Human Rights Commission, 2021). It has been a major impetus for the development of the Trans-Papuan Highway (Gaveau et al., 2021).*

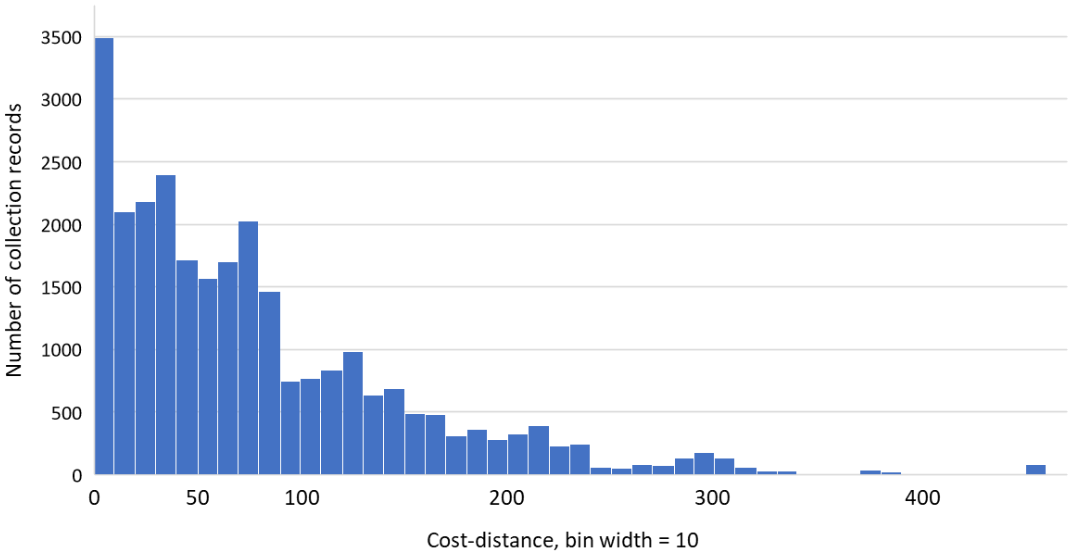


*Figure S3. Histogram of number of collection records by cost-distance, showing that few records are associated with high cost-distance. The collection sites with highest cost-distance values (450–460) were in the rainforests of the remote Louisiade Islands, to the SE of New Guinea.*

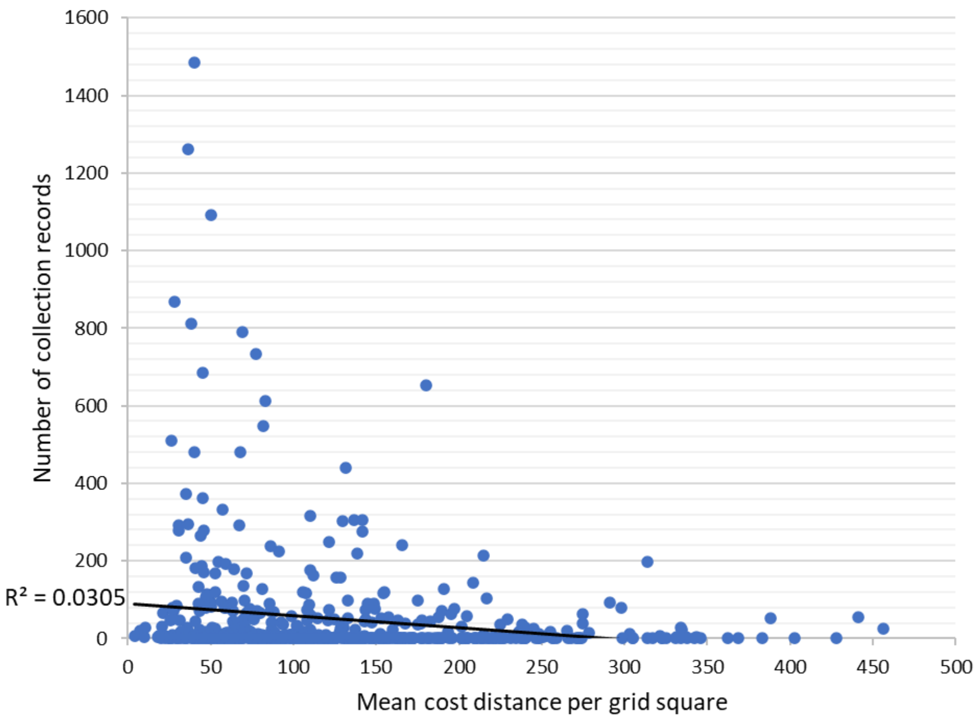


*Figure S4. Scatterplot of number of collection records (per 50km$^2$ grid square) against cost-distance, showing a non-significant relationship ($R^2$=0.03) due to a large number of grid squares with no collection records.*

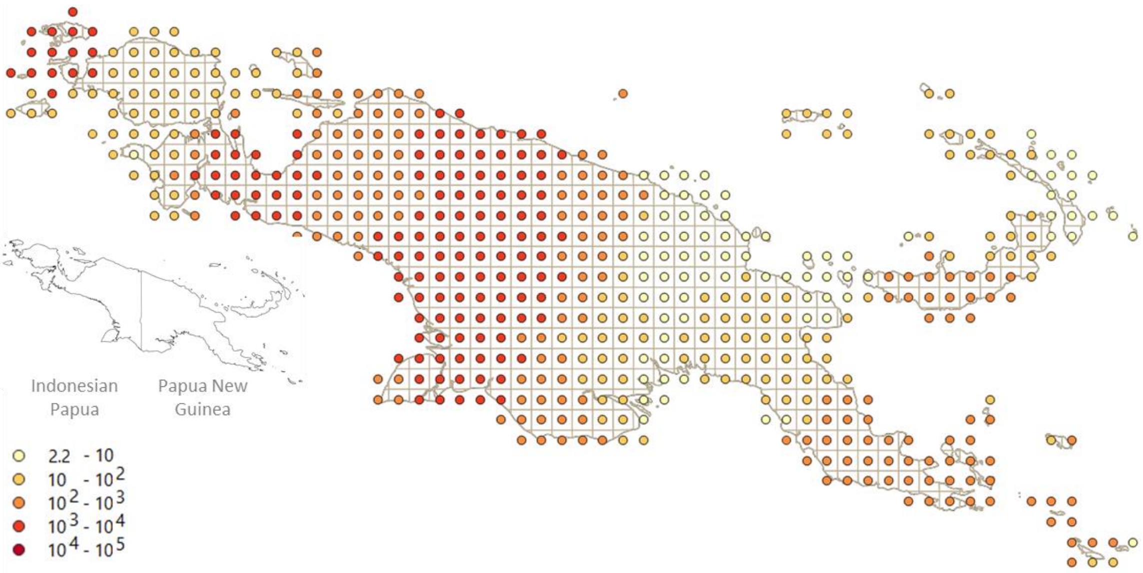


*Figure S5: Phylogenetic endemism (log scale, depicted by quintile). Predicted from HMSC model (5 PCA variables plus cost-distance). The bias-controlled model differs only slightly from the model shown here, which shows more even gradients across the region. Inset shows international boundary.*

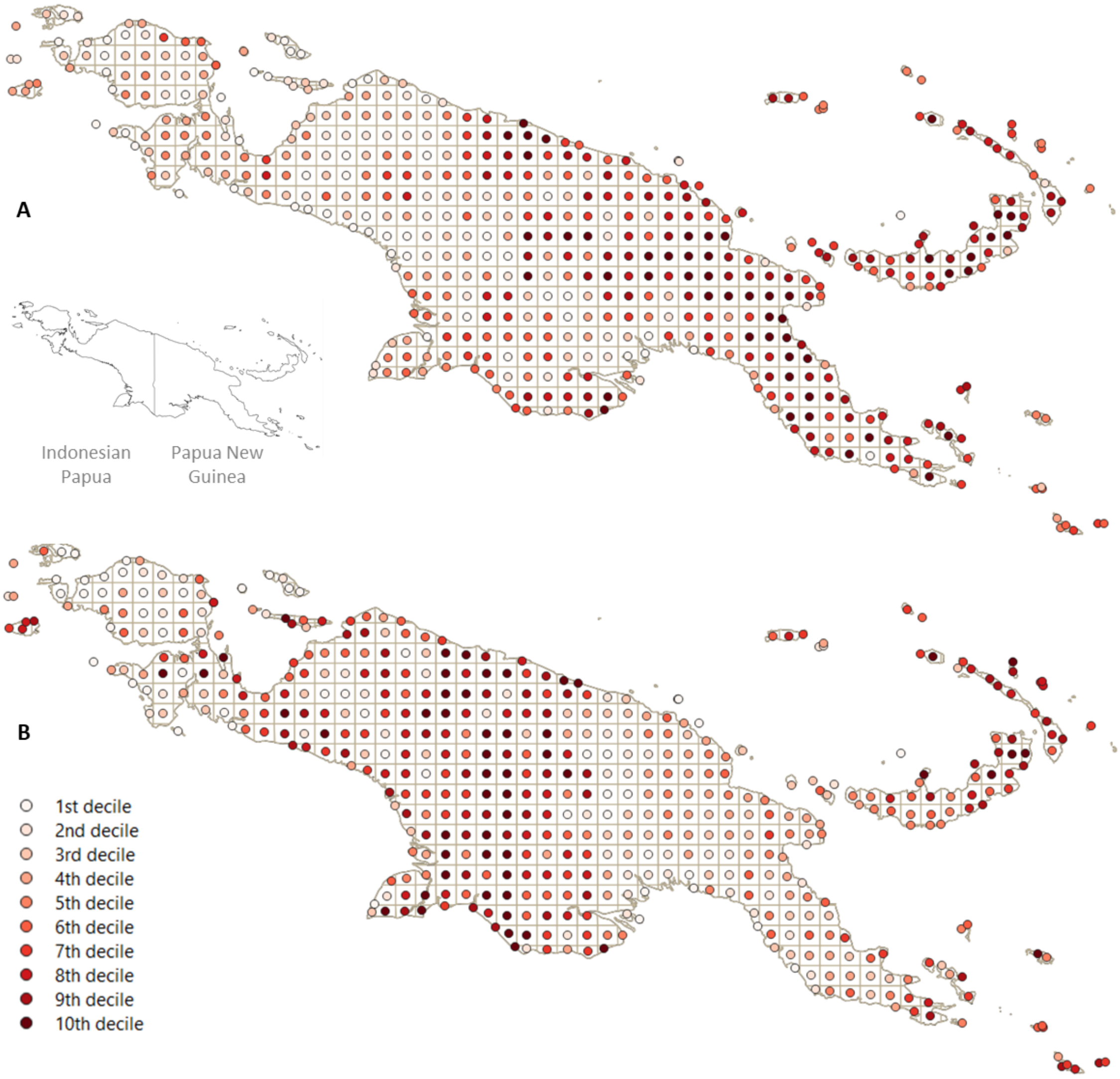


*Figure S6. Composite measures of predicted biodiversity combined with predicted deforestation (to 2025). A, calculated using biodiversity model with 5 PCA axes plus cost-distance variable. B, calculated using biodiversity model with cost-distance variable controlled. All ten deciles of the standardized measure are shown. Inset shows international boundary. The model shown in B (compared with that shown in A) suggests comparatively fewer hotspots of high biodiversity / high risk in the PNG central highlands, and a greater threat from plantations in Indonesian Papua (e.g. the MIFEE).*

*Table S1. Cost value assignment to land classification categories used during the preparation of the cost-distance map. The landsystems polygon map (Saxon & Sheppard, 2010) was rasterized, converting category values to pixel values, and pixel values were reassigned using QGIS/Raster analysis/Reclassify by table).*

| Land classification | Cost (0-10) |
|---|---|
| Roads | 1 |
| Lakes | 1 |
| Sea | 1 |
| Coastal plains with periodic brief flooding | 5 |
| Seasonally inundated coastal plains | 6 |
| Waterlogged plains | 7 |
| Permanently inundated coastal plains | 7 |
| Near-permanently inundated coastal plains | 7 |
| Montane and alpine wetlands | 7 |
| Flooded valley wetlands | 7 |
| Wetlands with back swamps | 9 |
| Peat swamps | 9 |
| Mangrove flats | 9 |
| All others (alluvial plains, coral islands, coastal flats, coastal plains, river valleys, foothills, ridges, peaks) | 3 |

*Table S2. Environmental variables used as Principal Component Analysis (PCA) inputs. Elevation map downloaded from United States Geographic Service (United States Geographic Service, 2021); MODIS, Moderate Resolution Imaging Spectroradiometer (Myneni, 2015); ISRIC, International Soil Reference and Information Centre (Poggio et al., 2021); FAO, FAO Soils Portal (Food and Agriculture Organization of the United Nations, 2021); ENVIREM, Environmental Rasters for Ecological Modelling (Title & Bemmels, 2018); Worldclim, www.worldclim.org (WorldClim, 2021).*

| Environmental variables |
|---|
| Elevation: corrected STRM data from 2000 |
| MODIS: Enhanced vegetation index |
| MODIS: Fraction of photosynthetically active radiation (400-700 nm) absorbed by green vegetation |
| MODIS: LAI (one-sided green leaf area per unit ground area in broadleaf canopies) |
| ISRIC: ph22o: Soil pH (5 variables: 0-5cm, 5-15cm, 15-30cm, 30-60cm, 60-100cm and 100-200cm below surface) |
| ISRIC: Bulk density of the fine earth fraction, cg/cm³ (5 variables: 0-5cm, 5-15cm, 15-30cm, 30-60cm, 60-100cm and 100-200cm below surface) |
| ISRIC: Cation Exchange Capacity of the soil, mmol(c)/kg (5 variables: 0-5cm, 5-15cm, 15-30cm, 30-60cm, 60-100cm and 100-200cm below surface) |
| ISRIC: Volumetric fraction of coarse fragments (> 2 mm), cm3/dm3 (vol%) (5 variables: 0-5cm, 5-15cm, 15-30cm, 30-60cm, 60-100cm and 100-200cm below surface) |
| ISRIC: Proportion of clay particles (< 0.002 mm) in the fine earth fraction, g/kg (5 variables: 0-5cm, 5-15cm, 15-30cm, 30-60cm, 60-100cm and 100-200cm below surface) |
| ISRIC: Total nitrogen (N), cg/kg, (5 variables: 0-5cm, 5-15cm, 15-30cm, 30-60cm, 60-100cm and 100-200cm below surface) |
| ISRIC: Proportion of sand particles (> 0.05 mm) in the fine earth fraction, g/kg (5 variables: 0-5cm, 5-15cm, 15-30cm, 30-60cm, 60-100cm and 100-200cm below surface) |
| ISRIC: Proportion of silt particles (≥ 0.002 mm and ≤ 0.05 mm) in the fine earth fraction, g/kg (5 variables: 0-5cm, 5-15cm, 15-30cm, 30-60cm, 60-100cm and 100-200cm below surface) |
| ISRIC: Soil organic carbon content in fine earth fraction, dg/kg (5 variables: 0-5cm, 5-15cm, 15-30cm, 30-60cm, 60-100cm and 100-200cm below surface) |
| FAO Soil Quality measure 1: Nutrient availability: Soil texture, soil organic carbon, soil pH, total exchangeable bases |
| FAO Soil Quality measure 2: Nutrient retention capacity (Soil Organic carbon, Soil texture, base saturation, cation exchange capacity of soil and of clay fraction) |
| FAO Soil Quality measure 3: Rooting conditions: Soil textures, bulk density, coarse fragments, vertic soil properties and soil phases affecting root penetration and soil depth and soil volume |
| FAO Soil Quality measure 4: Oxygen availability to roots: Soil drainage and soil phases affecting soil drainage |

| |
|---|
| FAO Soil Quality measure 5: Excess salts: Soil salinity, soil sodicity and soil phases influencing salt conditions |
| FAO Soil Quality measure 6: Calcium carbonate and gypsum |
| ENVIREM: Thornthwaite aridity index: Index of the degree of water deficit below water need |
| ENVIREM: climatic moisture index - a metric of relative wetness and aridity |
| ENVIREM: Emberger's pluviothermic quotient |
| ENVIREM: Count of the number of months with mean temp greater than 10℃ |
| ENVIREM: Continentality (mean temp of warmest month - mean temp of coldest month) |
| ENVIREM: Annual PET |
| ENVIREM: Mean monthly PET (potential evapotranspiration) of coldest quarter |
| ENVIREM: Mean monthly PET (potential evapotranspiration) of driest quarter |
| ENVIREM: Monthly variability in potential evapotranspiration |
| ENVIREM: Mean monthly PET of warmest quarter |
| ENVIREM: Mean monthly PET of wettest quarter |
| ENVIREM: Compensated thermicity index: sum of mean annual temp., min. temp. of coldest month, max. temp. of the coldest month, x 10 |
| ENVIREM: GrowingDegDays0 (sum of mean monthly temperature for months with mean temperature greater than 0℃ multiplied by number of days) |
| ENVIREM: GrowingDegDays5 (as above, temperature greater than 5℃) |
| ENVIREM: MaxTempColdestMonth |
| ENVIREM: MinTempWarmestMonth |
| ENVIREM: GIS-derived topographic wetness index |
| ENVIREM: GIS-derived terrain roughness index |
| Worldclim 2.1_bio: Max Temperature of Warmest Month |
| Worldclim 2.1_bio: Min Temperature of Coldest Month |
| Worldclim 2.1_bio: Mean Temperature of Warmest Quarter |
| Worldclim 2.1_bio: Mean Temperature of Coldest Quarter |
| Worldclim 2.1_bio: Mean Temperature of Wettest Quarter |
| Worldclim 2.1_bio: Mean Temperature of Driest Quarter |
| Worldclim 2.1_bio: Annual Mean Temperature |
| Worldclim 2.1_bio: Annual Precipitation |
| Worldclim 2.1_bio: Precipitation of Wettest Month |
| Worldclim 2.1_bio: Precipitation of Driest Month |
| Worldclim 2.1_bio: Precipitation of Wettest Quarter |
| Worldclim 2.1_bio: Precipitation of Driest Quarter |
| Worldclim 2.1_bio: Precipitation of Coldest Quarter |
| Worldclim 2.1_bio: Precipitation Seasonality (Coefficient of Variation) |
| Worldclim 2.1_bio: Precipitation of Warmest Quarter |
| Worldclim 2.1_bio: Mean Diurnal Temperature Range (mean of monthly max temp – monthly min temp) |
| Worldclim 2.1_bio: Isothermality (BIO2/BIO7) (×100) |
| Worldclim 2.1_bio: Temperature Seasonality (standard deviation ×100) |
| Worldclim 2.1_bio: Temperature Annual Range (BIO5-BIO6) |

#### 3. Supporting Information references


Asian Human Rights Commission (2021). *INDONESIA: MIFEE: The stealthy face of conflict in West Papua*. Retrieved August 27, 2021, from http://www.humanrights.asia/opinions/AHRC-ETC-022-2012

Food and Agriculture Organization of the United Nations. (2021). *Food and Agriculture Organization of the United Nations Soils Portal*. Retrieved August 26, 2021, from http://www.fao.org/soils-portal

Gaveau, D. L. A., Santos, L., Locatelli, B., Salim, M. A., Husnayaen, H., Meijaard, E., Heatubun, C., & Sheil, D. (2021). Forest loss in Indonesian New Guinea (2001–2019): Trends, drivers and outlook. *Biological Conservation*, *261*, 0–3. https://doi.org/10.1016/j.biocon.2021.109225

Microsoft Corporation. (2018). *Microsoft Excel* (No. 2107). https://office.microsoft.com/excel

Myneni, R. (2015). *MYD15A2H MODIS/Aqua Leaf Area Index/FPAR 8-Day L4 Global 500m SIN Grid V006 | NASA EOSDIS Land Processes DAAC*. https://doi.org/https://doi.org/10.5067/MODIS/MYD15A2H.006Poggio, L., De Sousa, L. M., Batjes, N. H., Heuvelink, G. B. M., Kempen, B., Ribeiro, E., & Rossiter, D. (2021). SoilGrids 2.0: Producing soil information for the globe with quantified spatial uncertainty. *Soil*, *7*(1), 217–240. https://doi.org/10.5194/soil-7-217-2021

QGIS.org. (2021). *QGIS Geographic Information System* (3.18 Zürich). QGIS Association. https://qgis.org

R Core Team. (2021). *R: A language and environment for statistical computing* (4.1). R Foundation for Statistical Computing, Vienna, Austria. https://www.r-project.org/

Saxon, E., & Sheppard, S. (2010). *Land Systems of Indonesia and New Guinea | Data Basin*. https://databasin.org/datasets/eb74fe29b6fb49d0a6831498b0121c99

The Mathworks Inc. (2021). *MATLAB* (MATLAB version 9.10.0.1613233 (R2021a)). The Mathworks, Inc.

Title, P. O., & Bemmels, J. B. (2018). ENVIREM: an expanded set of bioclimatic and topographic variables increases flexibility and improves performance of ecological niche modeling. *Ecography*, *41*(2), 291–307. https://doi.org/10.1111/ecog.02880

United States Geographic Service. (2021). *EROS Archive - Digital Elevation - Global Multi-resolution Terrain Elevation Data 2010 (GMTED 2010)*. Retrieved August 26, 2021, from https://www.usgs.gov/centers/eros/science/usgs-eros-archive-digital-elevation-global-multi-resolution-terrain-elevation?qt-science_center_objects=0#qt-science_center_objects

*WorldClim*. (2021). Retrieved August 26, 2021, from https://www.worldclim.org/
